# Deep Learning-Enabled, Detection of Rare Circulating Tumor Cell Clusters in Whole Blood Using Label-free, Flow Cytometry

**DOI:** 10.1101/2023.08.01.551485

**Authors:** Nilay Vora, Prashant Shekar, Michael Esmail, Abani Patra, Irene Georgakoudi

## Abstract

Metastatic tumors have poor prognoses for progression-free and overall survival for all cancer patients. Rare circulating tumor cells (CTCs) and rarer circulating tumor cell clusters (CTCCs) are potential biomarkers of metastatic growth, with CTCCs representing an increased risk factor for metastasis. Current detection platforms are optimized for *ex vivo* detection of CTCs only. Microfluidic chips and size exclusion methods have been proposed for CTCC detection; however, they lack *in vivo* utility and real-time monitoring capability. Confocal backscatter and fluorescence flow cytometry (BSFC) has been used for label-free detection of CTCCs in whole blood based on machine learning (ML) enabled peak classification. Here, we expand to a deep-learning (DL) -based, peak detection and classification model to detect CTCCs in whole blood data. We demonstrate that DL-based BSFC has a low false alarm rate of 0.78 events/min with a high Pearson correlation coefficient of 0.943 between detected events and expected events. DL-based BSFC of whole blood maintains a detection purity of 72% and a sensitivity of 35.3% for both homotypic and heterotypic CTCCs starting at a minimum size of two cells. We also demonstrate through artificial spiking studies that DL-based BSFC is sensitive to changes in the number of CTCCs present in the samples and does not add variability in detection beyond the expected variability from Poisson statistics. The performance established by DL-based BSFC motivates its use for *in vivo* detection of CTCCs. Further developments of label-free BSFC to enhance throughput could lead to critical applications in the clinical detection of CTCCs and *ex vivo* isolation of CTCC from whole blood with minimal disruption and processing steps.

## Introduction

Metastatic tumor growth is the leading cause of all cancer-related deaths^1^. During cancer progression, individual cells are observed to detach from the primary tumor and enter the bloodstream in a process known as intravasation^2^. Once in the bloodstream, these cells called circulating tumor cells (CTCs), can extravasate into distal organs, forming secondary tumors^3–6^. Multiple studies have correlated the dissemination of CTCs with poor prognosis and treatment resistance^7^.

During the metastatic cascade, CTCs and naturally occurring cells in blood can also form aggregates called CTC clusters (CTCCs)^1, 8–11^. CTCCs typically vary in size from as few as two cells to more than nine cells and are extremely rare, with less than four CTCCs being observed per 7.5 mL of blood^1, 7, 12^. While rare, CTCCs have gained significant attention due to their distinct characteristics and behaviors compared to individual CTCs. CTCC formation provides certain advantages to cancer cells, including increased survival rates in the bloodstream and enhanced ability to colonize distant tissues^1, 12^. The collective presence of multiple cancer cells within a cluster can provide protection against immune system attacks, promote resistance to therapies, and facilitate the formation of secondary tumors^1, 9, 10^.

While interest in CTC and CTCC detection and isolation has grown, the only FDA-approved technique to date is CellSearch^8, 13^. CellSearch is optimized for the enrichment, labeling, and detection of rare CTCs in whole blood with greater than 85% recovery^13, 14^. However, no conclusive data are available on the enrichment and detection of CTCCs by CellSearch, with only two studies listing anywhere from 0-53% enrichment efficiency^7, 13, 15, 16^.

Microfluidic and size-based approaches provide an epitope-independent technique for CTCC isolation^1, 7, 9, 17–21^. New isolation devices can provide up to 90% detection sensitivity for CTCCs in whole blood^21^. However, microfluidic devices depend on *ex vivo* blood processing of small volumes of blood compared to the total blood volume, leading to over or underestimation of CTCCs^12^. As liquid biopsy interrogation for CTCs and CTCCs has advanced, multiple groups have highlighted shifts in CTC dissemination due to hormonal changes during sleep cycles^22–26^. Further, the temporal selection of blood draws demonstrates high variability (order of magnitude or more) in CTC counts and consequently, CTCC counts, in as little as a few minutes^24, 25^. *Ex vivo* processing of blood samples in microfluidic channels is, therefore, likely to lead to poor correlation with prognosis.

*In vivo* flow cytometry (IVFC) provides a robust, highly sensitive and specific platform for CTCC detection continuously^5, 24, 27–33^. Fluorescence-based IVFCs (FIVFC) have been used to detect both rare CTCs and CTCCs; however, they are limited by the need for exogenous contrast agents^1, 5, 24, 27–30, 34^. Label-free IVFC (Lf-IVFC) systems utilize intrinsic contrast from CTCs and CTCCs, enabling wider clinical utility^31–33^. One such Lf-IVFC system, the photoacoustic flow cytometer (PAFC), has already demonstrated successful clinical detection of CTCCs *in vivo* in humans; however, the absorbance of melanoma cells is crucial in enabling detection of the CTCCs with this platform^31^. To expand the PAFC for broader use, photoacoustic contrast agents would need to be developed and approved by the FDA for *in vivo* use, limiting full clinical adoption.

A critical gap between broad CTCC detection and label-free techniques exists. To address this, our group has focused on developing label-free, backscatter flow cytometry (BSFC). BSFC monitors intrinsic light scattering and fluorescence to detect CTCCs^12, 35^. We have previously demonstrated using *in vitro* BSFC that CTCCs have unique light scattering signatures^35^, which can be used to detect and classify CTCCs in whole blood using machine-learning (ML) based algorithms^12^. However, exogenous fluorescence was used in these studies to identify CTCCs from non-CTCCs (NCs) events^12^. In this study, we aim to improve our ML model for fully label-free detection of CTCCs in whole blood and assess the clinical utility of BSFC for CTCC detection.

For the work described here, fresh rodent blood samples were spiked with green fluorescence protein- (GFP-) expressing CTCs and CTCCs. Light scatter and fluorescence data were collected using BSFC to design a peak detection and classification algorithm, herein referred to as the DeepPeak model. The model’s performance was assessed using the criteria proposed by Allard et al. (2004) for validation of the CellSearch platform^14^. Namely, we sought to answer two questions. First, what is the lowest number of CTCCs needed in blood to detect one CTCC? Second, what is the potential extent of variability at a theoretical level when measuring the reproducibility of rare events based on a random distribution^14^? We further assessed the error rate of BSFC on blood samples not expected to contain any CTCCs to determine the false alarm rate (FAR) of the DeepPeak model. Finally, we compared all relevant performance metrics reported using the key CTCC detection platforms. We demonstrate that the DeepPeak model with BSFC provides a clinically relevant, label-free CTCC detection platform with comparable performance to other CTCC detection platforms and unique potential to be extended to in vivo human studies.

## Experimental

### Sample Preparation

Blood samples were collected as previously described^12^. Briefly, 500 µL of blood from healthy, non-experimentally manipulated rats from other studies was collected via cardiac puncture immediately after CO_2_ euthanasia in K2EDTA-coated blood tubes^12^. All blood collections were performed in accordance with Tufts University Institutional Animal Care and Use Committee regulations (Protocol # M2022-132; formally M2019-158). All collected blood samples were processed within 24 hours of the blood draw.

CTCCs were introduced to the blood samples prior to flow data collection. MDA-MB-231 cells, a well-characterized human triple-negative metastatic breast cancer cell line, were used for all studies. CTCCs were generated using a previously established protocol^12, 18^. Briefly, GFP-associated MDA-MB-231 cells were grown on a 10 cm culture plate to 90% confluency. Following a wash step with phosphate buffer saline (Invitrogen), 1.5 mL of 0.25% trypsin (Gibco) was added to cleave the bonds between the cells and the plastic culture plate. As a result of the trypsin, natural aggregates (CTCCs) were observed to form (see Supplementary Fig. S1 online). Fully prepared media with serum was used to deactivate excess trypsin. Floating CTCCs were then carefully transferred for spiking into whole blood samples. Mechanical dissociation was expected to impact the size of CTCCs and the number of CTCs found in the sample; as such, it was critical to minimize introducing excessive forces during transfer and spiking steps.

During spiking, 100 µL of the mixture of CTCCs and CTCs were added to the blood tube. A tube rotator (VWR) was used to gently mix the CTCCs/CTCs into whole blood for 3-5 minutes. Once mixed, the samples were brought to the flow cytometer system for the collection of light scatter and fluorescence data. All studies conducted were approved by the Tufts University Institutional Biosafety Committee (Protocol # 2022-M71; formally 2020-M1) (Figure 1**a**).

**Figure 1:**
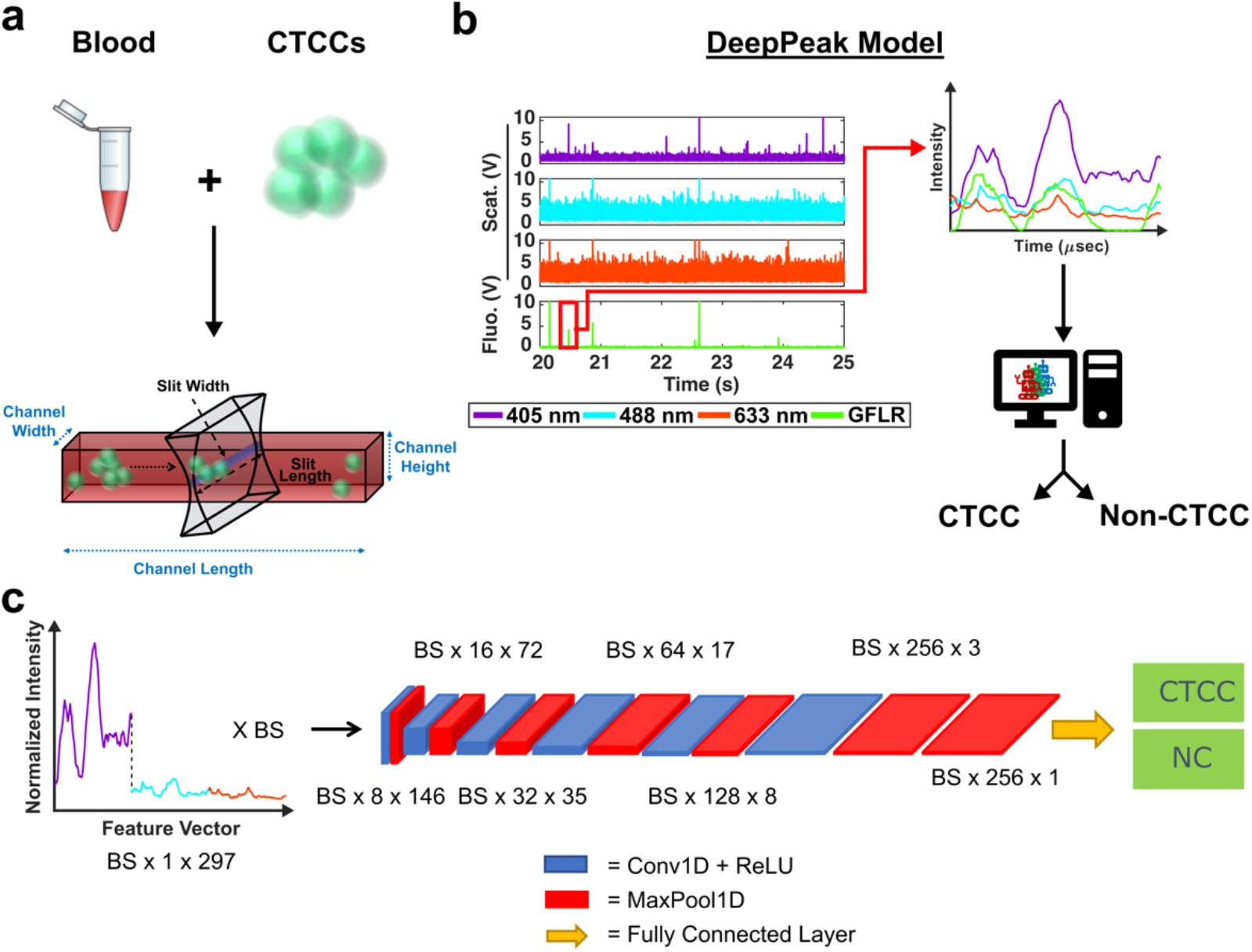
**(a)** Schematic of experimental design. Whole blood is collected for rodents and spiked with MDA-MB-231 breast cancer cell clusters. Data are collected through a microfluidic channel using a sharp illumination slit. **(b)** Data are processed using the DeepPeak model. Regions of interest (ROIs) are first selected using a new ROI detection algorithm before being passed to an ROI classification algorithm. **(c)** The classification algorithm utilizes a 1-D feature vector containing normalized scattering intensity from the three scattering wavelengths and a convolutional neural network (CNN) to classify CTCC peaks from Non-CTCC (NC) peaks.

### Flow Cytometer and Data Collection

The BSFC instrument has been previously described^12, 35^. A 405 nm, 488 nm, and 633 nm laser were used as excitation sources. Photomultiplier tubes (PMTs) were configured for the collection of light scattering from the three excitation lasers and red autofluorescence (670 ± 20 nm) and green exogenous fluorescence (525 ± 25 nm). Green exogenous fluorescence (GFLR) was used as a ground truth label for the location in the light scatter data corresponding to CTCC scattering.

Data were sampled at 60 kHz and stored using a data acquisition (NI-DAQ) unit (National Instruments; USB-6341). A custom LabVIEW (v18.0; National Instruments) project was written to read data output from the NI-DAQ and save it as a comma-separated values (CSV) file. A wrapper function was written in MATLAB to read the CSV files and store the data into smaller 1.5-minute-long data segment. Each segment was then processed for CTCC detection by the DeepPeak model (Figure 1**b**), which was composed of a region-of-interest (ROI) Detection (Figure 2) and ROI Classification Algorithm (Figure 1**c**).

**Figure 2:**
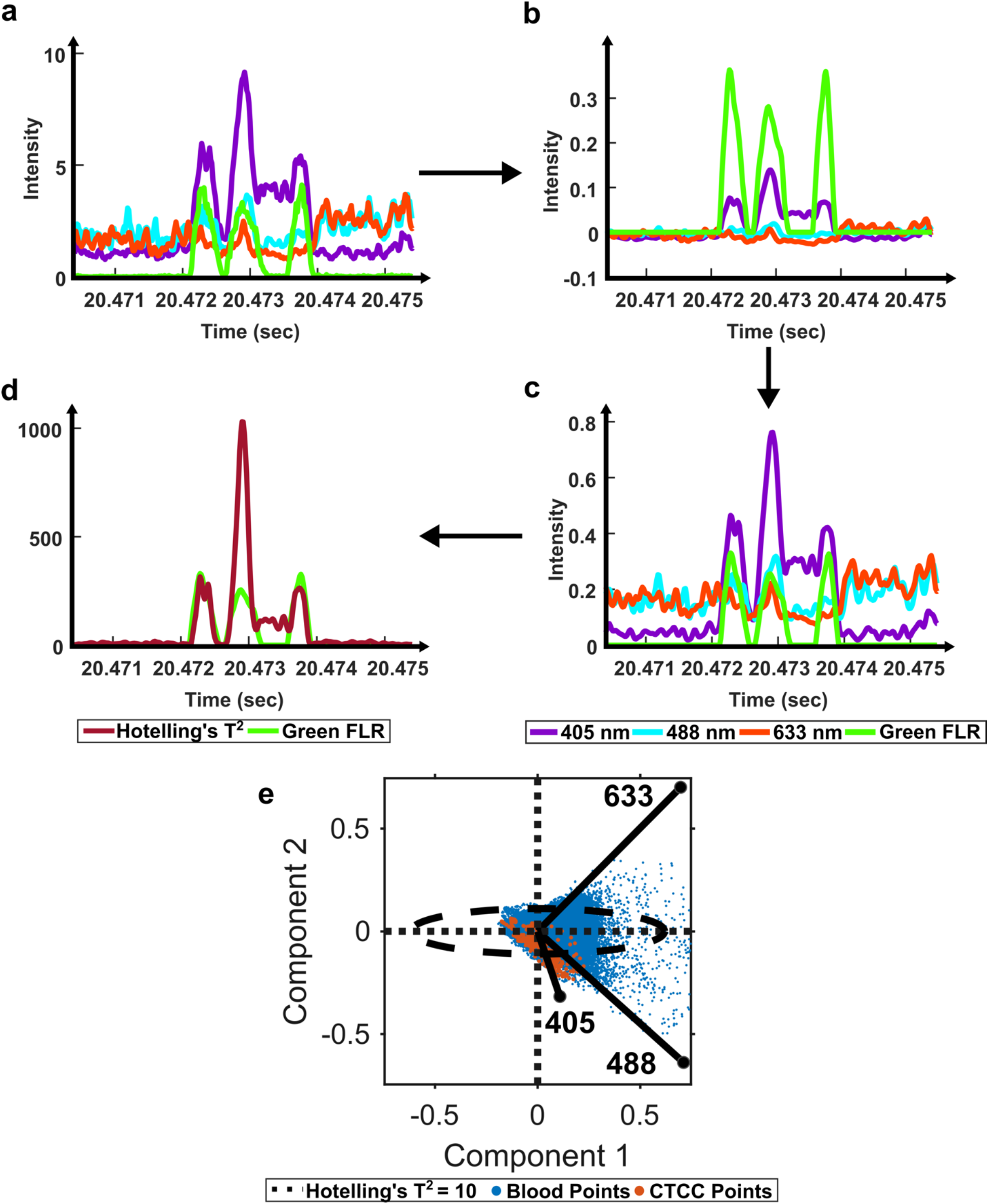
Schematic of the ROI Detection Algorithm. **(a)** Raw data is loaded in and **(b)** filtered using a second-order Butterworth filter. **(c)** Filtered data is normalized between 0 and 1 using the maximum intensity. **(d)** PCA and Hotelling’s T^2^ test are used to calculate the distance of each observation from the centroid, generating a new cumulative scattering trace. **(e)** Loadings for two principal components and the corresponding data are shown, along with the threshold used for peak detection. Points outside the oval represent strong outliers and are part of a suspected CTCC peak.

### ROI Detection Algorithm

To detect ROIs, data segments were processed one at a time. Baseline variability was assessed to determine whether blood clots existed in the data. A blood clot was found to be present if the standard deviation of the cumulative scattering signal was greater than 1.75. A threshold of 1.75 was selected based on the inspection of data without clots. Blood clots were characterized by the rise in the baseline signal due to increased background scattering signal. If variability greater than 1.75 was detected in the segment, a 500-point moving average of the signal was calculated for all points to find the average baseline signal. This baseline was removed from all points in the segment to exclude the shift in background scattering intensity from the blood clot (zero-mean data). To preserve the scattering signal’s positive values, the mean intensity of the entire signal was added back to the zero-mean data. This process was repeated up to three times or until the standard deviation of the scattering signal fell below 1.75, whichever came first. A maximum of three was selected to prevent an infinite loop on noisier data.

Once cleaned, we proceeded with previously described standard preprocessing steps^12^. Specifically, a second-order Butterworth filter was used to remove high-frequency noise and normalize the baseline signal (50-10000 Hz) (Figure 2**b**). Then, the filtered light scatter signal was normalized for differences in power measurements from day to day (Figure 2**c**). Normalized and filtered data were used for all ROI detection steps.

To extract ROIs, anomaly detection methods were implemented. Principal component analysis (PCA) has been used for various applications to extract features from inter-correlated data^36, 37^. PCA extracts the most important features from multivariate data and reduces the dimensionality to compress the data^36^. In the case of anomaly detection, outliers, like scattering from CTCCs and CTCs, were expected to contribute the most to data variability^38, 39^. Using this principle, we first used PCA to reduce the dimensionality of our light scatter dataset. We then assessed the anomalies in the dataset using a statistical test called Hotelling’s T^2^ test (Figure 2**d**)^37–39^. Hotelling’s T^2^ test measures the squared Mahalanobis distance of each point from the centroid of the principal components^38, 39^. Outliers were characterized by larger magnitudes, while inliers featured little to no magnitude. As Hotelling’s T^2^ values were calculated at each point in the dataset, the data were reformatted to measure outlier probability over time (Figure 2).

Previously described ROI detection algorithms were then used to locate ROIs in the outlier time-series dataset^12^. Briefly, the built-in MATLAB (R2021b, Natick, MA) function findpeaks.m was used to find local maximums in the dataset. A simple intensity threshold of ten was set based on experimentation to maximize initial detection sensitivity and purity (See Supplementary Fig. S2 online). Locations where the outlier signal crossed the intensity threshold were used to extract the peak event ranges. Each event range was inspected to remove extra peaks within a range as we sought to label the entire ROI as a single cluster event. Peak characteristics such as full-width-at-half-max (FWHM), location, and intensity were recorded for all events. During data visualization, we observed peaks with narrower than expected FWHM values due to lower-intensity shoulder peaks (see Supplementary Fig. S3 online). To correct for differences in the height of shoulder peaks during FWHM measure, a geometric height equalization algorithm was implemented^40^. In this equalization algorithm, all local maxima were rescaled to one, and points in-between were scaled by a fitted line from peak to peak. Once peaks were equalized, standard FWHM measurements on the equalized signal were possible. A spreadsheet containing peak characteristics was saved at the end of this step for peaks found in both the scattering and fluorescence acquisition channels. The green fluorescence channel was used as a ground-truth label for CTCCs, while the light scattering data was used for label-free detection of the CTCCs.

As these studies aimed to demonstrate label-free detection of CTCCs in whole blood, peaks from single cells were removed using a peak width threshold^12^. To calculate the threshold, the estimated size for a large CTC or white blood cell (12-15 μm) was used in combination with the flow speed (55.6 mm/sec) to calculate the maximum time it would take for a large single event to cross the illumination slit. As the sample rate was 60,000 samples per second, we anticipated a single cell would measure 21-22 points in width. To calculate the corresponding FWHM, we multiplied the full peak width by 0.75, which represented a conservative measure of the relationship between FWHM and event width. Therefore, the calculated threshold for multicellular events was set to 17 points. Peaks less than 17 points in FWHM were removed, with the remaining peaks selected as ROIs.

### ROI Classification Algorithm

Once ROIs were identified, feature vectors were generated for the ROI classification algorithm. Feature vectors were designed similarly to previous work^12^. Briefly, raw data was loaded individually and normalized by subtracting the mean signal and dividing it by the standard deviation (Zero-Mean Normalization). The normalized data were then parsed based on identified ROIs from the ROI detection algorithm. Data from the 405 nm, 488 nm, and 633 nm channels were collected in a window of ±49 points from the peak location. A window size of ±49 points was selected to ensure that large and small clusters would be fully included in the feature vector. The three sets of 99 data points were concatenated to generate a single 297-point feature vector (405nm channel = features 1-99, 488 nm channel = features 100-198, and 633 nm channel = features 199-297). To generate a label, ROIs identified in the green fluorescence channel were cross-referenced with ROIs from the scattering channel. If a matching peak was found in the green fluorescence channel and the scattering channel, the event was labeled as a CTCC event; if the event was only found in the scattering channel, it was labeled as a Non-CTCC (NC) event. A total of 34 independent days of experimental data were formatted for classification. 32% of the data (11 days) were set aside as a test set. The remaining 68% were used to train and validate the ROI classification model. During training, 21% of the training set (5 days) was used for validation, with the remaining 18 days used for training. A total of five training-validation folds were used to verify model performance.

The Tufts High Performance Cluster was used for all ROI classification algorithm training. A single, eight-core CPU with a 40-gigabyte Nvidia Tesla A100 GPU card was used for all training and evaluation. The classification algorithm utilized a convolutional neural network (CNN) to classify NC events from CTCC events accurately. The CNN architecture was based on prior work by Melnikov et al. (2020), which examined peak detection in noisy liquid chromatography−mass spectrometry (LC−MS) data^41^. The designed CNN featured six convolutional + max pooling layers followed by an additional max pooling layer and a fully connected layer (Figure 1**c**).

The classification algorithm was implemented using PyTorch in an anaconda environment^42, 43^. A starting learning rate of 1 x 10^-3^ was used with an Adam optimizer^44^. During training, the maximum number of epochs was set to 15 with an early stop condition if performance failed to improve after seven epochs. As class imbalance was expected to be significant, a weighted binary cross-entropy (BCE) loss function was combined with the Focal Tversky Loss function^45, 46^.

BCE loss is a robust loss function for equally balanced datasets; however, in the case of high-class imbalance, models learn little from the misclassification of the minority class. Weighted BCE loss attempts to improve application on imbalanced datasets by increasing the penalty on minority class misclassification. However, weighted BCE may not perform well on highly-imbalanced datasets^46^. Focal Tversky Loss (FTL) was designed for use on highly-imbalanced datasets^45^. FTL enables flexibility in false negative (FN) and false positive (FP) detection based on hyperparameters controlling the acceptable limits of FNs and FPs. However, FTL can be unstable in learning based on parameter selection. To account for this, we combined BCE loss with FTL to stabilize learning while promoting accurate classification of a largely imbalanced dataset.

To further improve the model’s performance, we used an ensemble of CNNs to improve detection purity. Each model was independently trained based on the output from the previous model. For example, model one was trained until performance stabilized, after which all FPs, FNs, and true positive (TP) events were separated from the events the CNN accurately classified as NC peaks (True negatives; TN). The isolated FP + FN + TP events were then inputted into the second CNN as the training set. This process was repeated for ten networks. The assumption was that each successive CNN would learn new boundaries to separate hard-to-discern NC and CTCC peaks. After training, the test set was evaluated through all ten networks. During evaluation, only the FP and TP events were passed as inputs into the subsequent network. Performance was logged after each network. The number of networks used was selected after the performance was observed to stagnate. The final classification algorithm’s performance was assessed after all ten networks had evaluated the test data set.

### Metrics

To determine the performance of the DeepPeak model, six metrics of performance were examined: Purity (also referred to as precision), Sensitivity, Specificity, FAR, F_1_ Score, and Pearson Correlation Coefficient (PCC) between the predicted number of events and the number of spiked events present (calculated based on green fluorescence signal). These metrics were defined as follows:

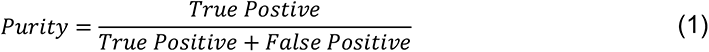

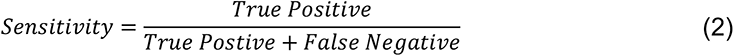

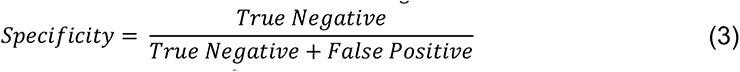

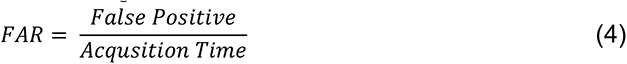

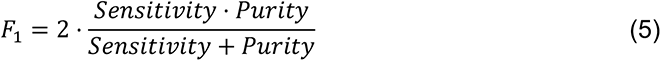

Metrics were selected to compare the DeepPeak model’s performance against other CTCC detection platforms. Performance targets were not available for all metrics from other platforms, but we present a complete list of all reported performance metrics below.

### Statistics

Poisson statistics have long been used to describe randomly distributed objects in a given volume and are frequently used to describe the detection of CTCs in liquid biopsies^14, 47^. Based on Poisson statistics, to detect an average of *x* events at a probability of detection (*p)*, a minimum of *n* events would be needed in the sample (equation 6). In this study, we assessed the minimum volume of blood needed to detect a minimum of 1 CTCC based on a Poisson distribution.

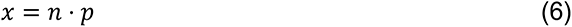

To calculate the necessary interrogation volume of blood, we first determined the minimum number of CTCCs needed in a sample to detect 1 CTCC based on the DeepPeak model’s sensitivity. Then, we estimated the blood volume necessary to detect a single CTCC based on the average concentration of CTCCs in patient blood. Concentrations of CTCCs in patient blood varied considerably from study to study, with some studies listing concentrations as low as 0.44 CTCCs/mL of blood^7^ and as high as 10 CTCCs/mL of blood^48, 49^. For the studies listed here, we assumed an average concentration of 0.4-0.5 CTCCs/mL of blood.

Additionally, to assess the reliability of measurements using the DeepPeak model, the coefficient of variability (CV) was calculated for multiple spiking ratios. CV was used as an alternative measure of standard deviation to assess variability in measurements without including mean^50^. Standard deviation shifts proportionally according to the mean number of events in a sample; however, these shifts make comparing the variability between different concentrations difficult^50^. CV accounts for differences in concentration by removing the mean and standardizing the variation.

CV was used in this study to assess the variability of the DeepPeak model when a restricted number of CTCCs were provided from various days of experimental measurements. Artificial spiking was completed in place of manually spiking a specific number of CTCCs into whole blood samples. CTCCs were observed to break apart during the isolation and spiking steps needed for manual spiking. This necessitated alternative methods of spiking such as artificial spiking. To simulate manual spiking, a set number of CTCC peak events were isolated from BSFC datasets along with all NC peak events leading up to the set CTCC count. For example, if 50 CTCCs were desired and the 50^th^ CTCC was found 30 minutes after collection started, all NC peaks found within the first 30 minutes of data were isolated along with the 50 CTCC peaks. CV values were calculated for spiked ratios of 5, 10, 30, 50, and 100 CTCCs.

To determine if the DeepPeak model added additional variation to the inherent variation of counting random events due to Poisson statistics, we calculated the theoretical variability (equation 7) for the five spiked CTCC concentrations and compared it to the observed %CV. As the volume of blood scatter peaks (NC peaks) varied based on the amount of time needed to detect the desired number of CTCCs, we estimated the variability in volume and accounted for this in our calculation of theoretical %CV through the sum of variance (equation 8).

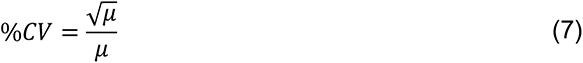

where *μ* is the average number of events in a sample.

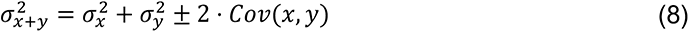

Where *x* is the measurement of theoretical variability and *y* is the measurement of volume variability.

## Results

### Assessment of the ROI Detection Algorithm

The first step in the DeepPeak model was to define potential ROIs within the time series data traces. In prior studies, we determined ROIs using an intensity threshold based on the variance in the baseline signal^12, 35^. Cumulative scattering intensity was interrogated using a built-in MATLAB® function findpeaks.m for all local maxima. Once all local maxima were identified, any maximum with an amplitude less than three to five times the standard deviation of the ninety seconds data segment was removed^12, 35^. The basis of this algorithm was to remove noise from the detectors and weaker scattering events. When implemented on datasets of depleted blood data (containing only white blood cells), the algorithm provided highly sensitive and specific detection of CTCCs^35^. However, when applied to CTCC detection in whole blood, ROI detection sensitivity fell to 43.3% with a detection purity of 0.2%.

It was assumed that whole blood scattering and absorption properties, originating particularly from red blood cells (RBCs) and plasma, contributed to the loss in sensitive and specific detection of CTCCs. RBCs and plasma account for up to 99% of whole blood samples and most of blood’s absorption and scattering properties^51^. Poorly defined peak characteristics due to absorption and increased baseline scattering from RBCs and plasma led to reduced detection signal-to-noise ratio (SNR) for CTCCs. As such, sensitive or precise detection of CTCCs using our standard threshold-based ROI algorithm was not possible.

To account for the reduced performance, an anomaly ROI detection algorithm was written using PCA and Hotelling’s T^2^ test to redefine how cumulative light scattering data were calculated. Whole blood backscatter intensity contributed heavily to our baseline signal and was a majority of the detected signal. It was therefore assumed that blood cell scattering would have lower Hotelling’s T^2^ metric values. Conversely, significant changes in scattering intensity from the baseline signal would have high Hotelling’s T^2^ metric values. As CTCCs have lower absorption and different scattering properties compared to RBCs, it was assumed that high Hotelling’s T^2^ metric values were from CTCCs. Based on this principle, we applied an empirical threshold of ten to detect outlier locations in the Hotelling’s T^2^ metric time trace (Figure 2**e**). Hotelling’s T^2^ values greater than ten were identified as outliers (outside of the ellipse), while those inside the ellipse were considered inliers and removed. The selection of ROIs based on the outlier points demonstrated an improvement in detection sensitivity from 43.4% to 85.1% and detection purity from 0.2% to 2%. This suggested that PCA and Hotelling’s T^2^ test could be used to extract CTCC ROIs.

### Assessment of the ROI Classification Algorithm

The second portion of the DeepPeak model was the classification algorithm. Despite the improvement in sensitivity and purity, 2% detection purity was far below the desired performance for ultimately *in vivo* clinical CTCC detection. To improve detection purity, we utilized a CNN-based classification algorithm. The classification algorithm included 10 CNNs ensembled to enhance the detection of rare cellular events in whole blood. The ensemble procedure was consistent with prior implementations completed by our group^12^. Peaks were labeled before classification based on the width of the scattering peak and ground truth (GFLR) signal. If the peak was greater than 17 points in FWHM in the cumulative light scatter trace and featured GFLR signal or was greater than 17 points in FWHM in the GFLR channel, the peak was considered a CTCC. Sample CTCC peaks are shown in Figure 3**a**. Classification performance was assessed using Purity, Specificity, Sensitivity, Accuracy, and Pearson Correlation Coefficient. *K-*fold validation was used to verify the reproducibility of the classification on varying validation sets with a *k*=5. On an independent test set, we observed approximately 69.0% detection purity, 98.7% specificity, 43.8% sensitivity, 95.5% accuracy, and r = 0.93 correlation between detected events and spiked events (Figure 3**b**). Across the entire dataset, including training and validation data, performance was stable with approximately 72.5% detection purity, 98.6% specificity, 60.5% sensitivity, 96.5% accuracy, and r = 0.94 correlation between detected events and actual events (Figure 3**b**). Confusion matrices for the test set and the full dataset for one of the folds are shown in Figure 3**b**.

**Figure 3:**
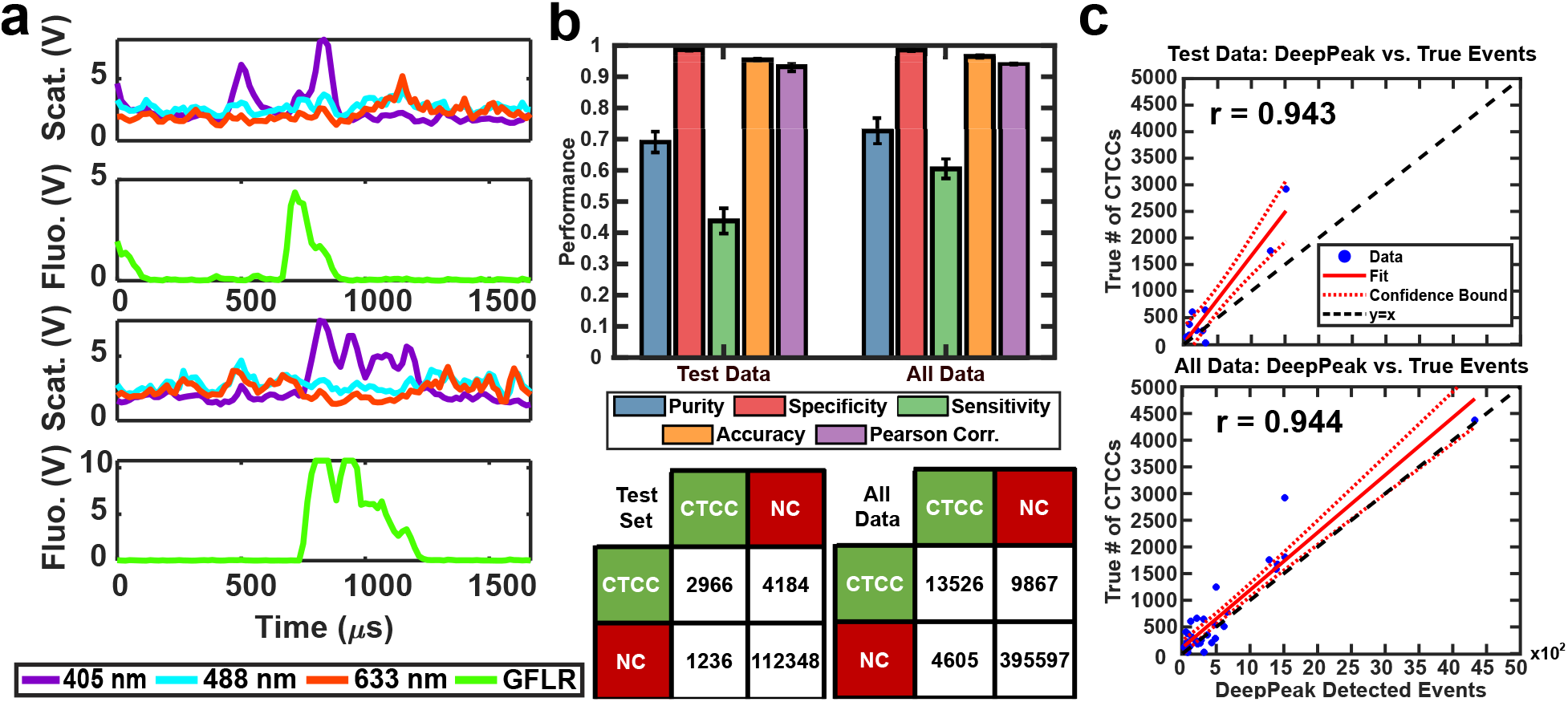
**(a)** Sample CTCC peaks. Peaks are defined as CTCCs based on the GFLR signal and FWHM. **(b)** Assessment of 5-fold validation on an independent test set and the full dataset (All Data). Confusion matrices are shown for the test set and full dataset. **(c)** Correlation plots for true # CTCCs compared to the number of events detected by the DeepPeak model.

A challenge in rare event detection was the large class imbalance. Consequently, while the classification algorithm improved detection purity, the low TP rate led to decreased sensitivity. Pearson correlation coefficient (PCC) was used to assess how well the detected events correlated with the anticipated CTCC count. We observed that the test set and full dataset detected event counts correlated highly with the expected number of events (Figure 3**c**). To further assess the impact of outliers in the linear fit, we refit the lower 30% of peak counts in the full dataset. PCC of the full dataset fell from 0.94 to 0.88 (data not shown), suggesting that while the outliers impacted our fit, the detected events were still well correlated with the actual event counts.

### Assessment of the DeepPeak Model on Unspiked Blood Data

For *in vivo* clinical utility, it was important for the DeepPeak model to minimize the number of FP events reported when no CTCCs were present, i.e., the false alarm rate (FAR). To assess the FAR of the DeepPeak model, unspiked blood samples from control animals were flowed for up to 60 minutes. Collected data were processed using an identical ROI detection algorithm as the spiked blood samples. Selected ROIs were then classified using the trained ROI classification algorithm. 175 minutes of data from five experimental days were used for FAR assessment. A total of 137 FP events were detected in the negative control blood dataset (Figure 4**a**). Detected FP events mimicked many of the characteristics of CTCCs in the light scatter channel (Figure 3**a** and Figure 4**b**). Interestingly, some FPs displayed weak autofluorescence, although the source of the autofluorescence was not confirmed in this study (Figure 4**b**). Based on the detected number of events in the control blood samples and time of collection, the FAR was estimated to be 0.78 events/min.

**Figure 4:**
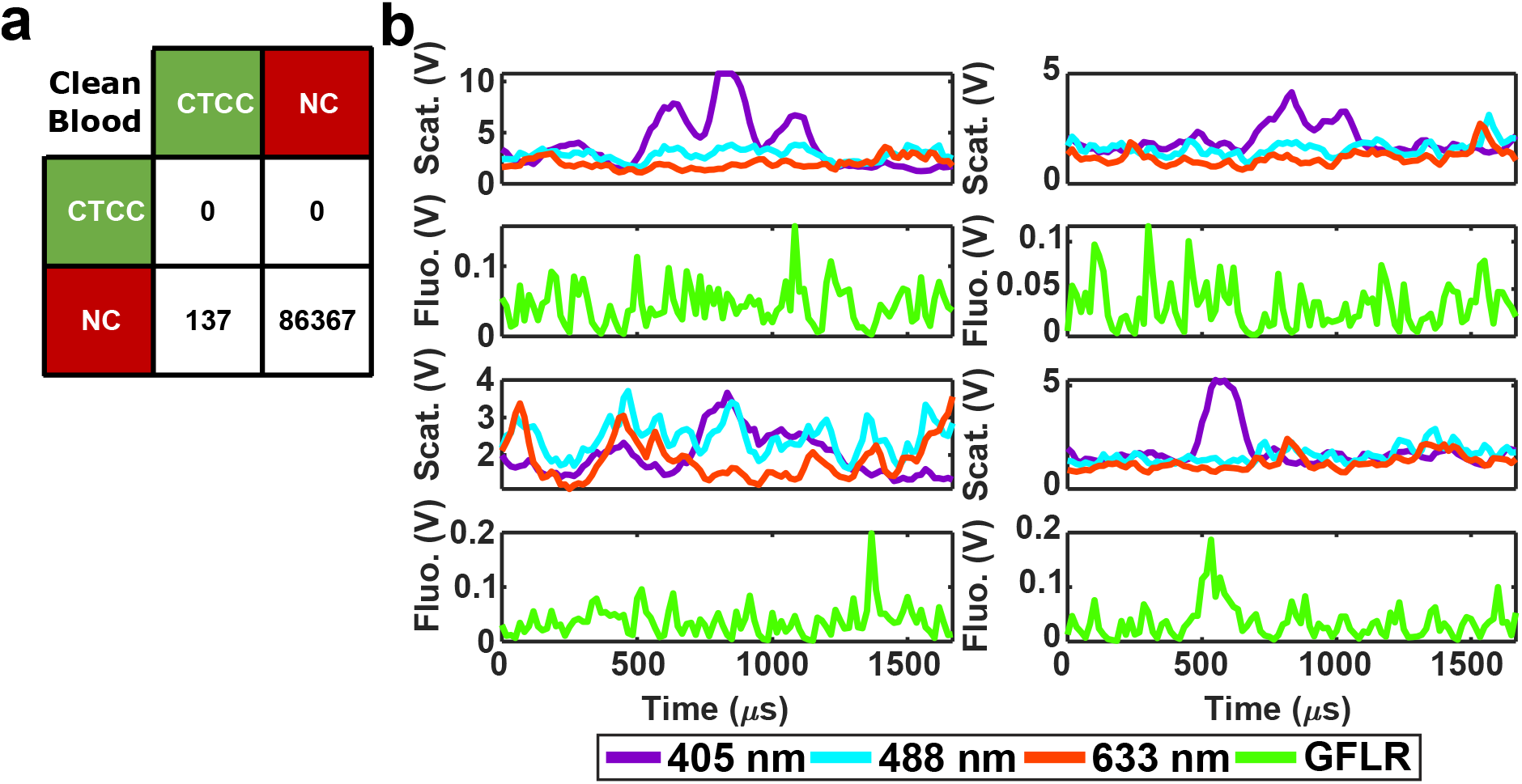
**(a)** Confusion matrix for events in negative control blood data after classification by the DeepPeak model. **(b)** Four examples of misclassified peaks by the DeepPeak model. Paired images show light scattering and fluorescence signals. Autofluorescence is observed in some FP events, such as the bottom right event.

### Assessment of the DeepPeak Model Consistency in Spiked Samples

Spiking CTCCs into blood has frequently been used to assess device performance in cell sorting and flow cytometry studies. However, to simulate *in vivo* concentrations of CTCCs, controlled spiking studies were needed to validate how sensitive the DeepPeak model was to changes in CTCC concentration and the limit of detection for the DeepPeak model. Further, replication of spiked CTCC concentrations enabled us to validate the detection consistency of the DeepPeak model. Five concentrations were selected: 5, 10, 30, 50, and 100 CTCCs. Formatted datasets containing the specified number of CTCCs and an assortment of NC peaks were evaluated by the trained model. The DeepPeak model demonstrated high sensitivity to changes in the concentration of CTCCs (r = 0.996) (Figure 5**a**). At a minimum, the

**Figure 5:**
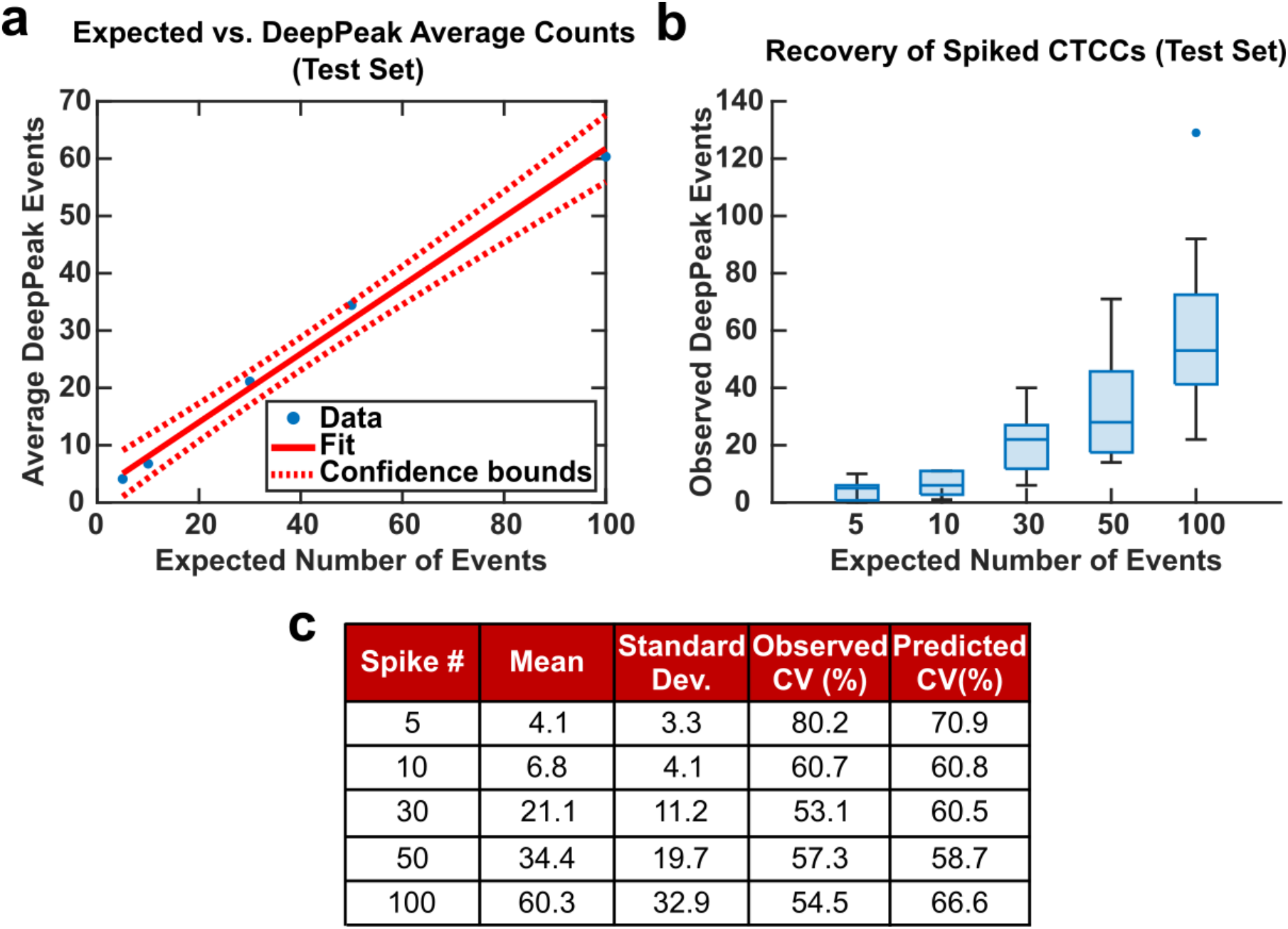
**(a)** Linear fit for controlled spiking study between the number of events detected by the DeepPeak model and the expected number of events. **(b)** Box plot for the various spike concentration compared to the number of events detected by the DeepPeak model. Variability is observed to increase with the number of events spiked in. **(c)** Summary table for assessing sources of variability in spiked cell study.

DeepPeak model recovered 60.3% of the expected counts. To assess the variability in detection, theoretical %CV and observed %CV were calculated for each spiked concentration. Despite the variance increasing with the number of expected events (Figure 5**b**), %CV decreased for all spiked concentrations outside of the 100 CTCC spiked concentration (Figure 5**c**). More significantly, observed %CV values were similar to the predicted (theoretical) %CV values. This implied that variability resulted from rare event detection based on Poisson statistics and no variability was added by the DeepPeak model. Overall, this suggested that the DeepPeak model could measure changes in and reliably assess CTCC concentrations.

### Net DeepPeak Model Performance

To assess the overall performance of the DeepPeak model, initial ROI detection sensitivity and specificity were combined with the ROI classification performance. For all performance metric analyses, we used weights from an ensemble of models trained on a specific data fold. Net sensitivity was calculated by multiplying the ROI detection algorithm sensitivity with the ROI classification algorithm sensitivity for the test set. Based on this formula, the observed net sensitivity for CTCC events was 35.3% (85.1% * 41.5%). To assess the net specificity, the total number of NC scattering events labeled by both the ROI detection and classification algorithms were summed together (TN_total_ = TN_detect._ + TN_class._) and compared to the total number of NC scattering events (FN + TN_total_). In total, the DeepPeak model demonstrated a net specificity of 99.97%.

Additional metrics considered included the FAR and F_1_ score. Minimizing FAR was considered important in preventing the misidentification of rare events when handling clinical samples. The F_1_ score was calculated to determine how well the model performed as a harmonic mean of sensitivity and purity. A high F_1_ score would indicate that both sensitivity and purity were high; however, a low F_1_ score could indicate that the performance favored only high sensitivity/high purity or had low sensitivity and purity. For clinical use, it was important for the DeepPeak model to be both sensitive to CTCCs and to minimize the number of FPs detected (maximize purity), as such, achieving high F_1_ scores was desirable. Owing to the high detection purity/specificity of the DeepPeak model, less than one FP event per minute of data collection (FAR = 0.78/min) was measured (Figure 4). Further, based on our detection purity and sensitivity (72% and 35%, respectively), the F_1_ score was approximately 0.474.

### Discerning the Clinical Utility of BSFC and the DeepPeak Model

To demonstrate the potential clinical value of BSFC and the DeepPeak model, we explored the final DeepPeak model performance metrics in the context of clinical use. Allard et al. (2004), in their characterization of the CellSearch platform for CTC detection, provided a blueprint for contextualizing the clinical utility of a rare event detection platform^14^. Two questions were proposed in the study to quantify the performance of rare event detection platforms.

The first question addressed the minimum number of CTCCs needed in a blood sample to detect a single CTCC. As the DeepPeak model’s net sensitivity was ∼35.3%, to detect a single CTCC, we would need ∼3 CTCCs (1/0.353) to be present within the sample. Using a relatively low estimated concentration of CTCCs in whole blood (0.4-0.5 CTCCs/mL of blood), we estimated that 5-7 mL of blood would need to be processed to detect a single CTCC. Accounting for our current throughput (3 μL/min), this would require between 27-39 hours of blood processing.

The second question proposed by Allard et al. (2004) was to determine the extent of variability present in measuring rare events reproducibly based on a random distribution. In artificially spiked samples, we observed variability in measurements that were consistent with theoretical values of variability at varying spike concentrations (Figure 5). Our results suggested that BSFC with the DeepPeak model did not increase the variation in CTCC measurements beyond the inherent variation in a random distribution. As the only source of variation originated from Poisson statistics for counting rare events, we determined that BSFC with the DeepPeak model could reliably detect rare CTCC events.

To understand the performance of our model in the context of broader CTCC detection, we compared our performance values against three flow cytometer systems, CellSearch, and two microfluidic platforms that have been used for CTCC detection (Table 1)^7, 16, 18, 19, 31, 52, 53^. Comparisons between different platforms were challenging due to numerous studies reporting a mixture of performance metrics from varying event types. However, we sought to identify performance targets based on the listed metrics in literature to the best of our ability.

**Table 1:** Summary of all performance metrics available from various CTCC detection platforms. Metrics are calculated using Equations 1-5. For clarity in the comparison of metrics, the type of cell events included in performance calculation is also indicated. Label-free systems are bolded.

| System | Purity | Sensitivity | Specificity | False Alarm Rate | F<sub>1</sub> Score | PCC | Event Inclusion |
| --- | --- | --- | --- | --- | --- | --- | --- |
| **PAFC**<sup>31</sup> | - | 62 ± 18% | 94.7% | - | - | - | CTC/CTCC |
| DiFC<sup>5,53</sup> | No comparable metrics discussed in paper |  |  | 0.017 per minute | - | 0.906 | CTC/CTCC |
| VIFFI FC<sup>52</sup> | - | 29% | 99.9984% | - | - | - | CTC/CTCC |
| **DLD Chip**<sup>18</sup> | - | 66.7 ± 6.4% | - | - | - | - | CTCC |
| **NISA-XL Chip**<sup>19</sup> | 5.5% | 84% | - | - | 0.1 | - | CTCC |
| CellSearch<sup>7,16</sup> | - | 0-53% | - | - | - | - | CTCC |
| **DeepPeak (Our Model)** | 72.0% | 35.3% | 99.97% | 0.78 per minute | 0.474 | 0.943 | CTCC |

In our results, we observed improved detection purity compared to the non-equilibrium inertial separation array-extra-large (NISA-XL) chip for CTCCs. However, sensitivity trailed both the deterministic lateral displacement (DLD) chip (CTCCs only), NISA-XL chip (CTCCs only), and PAFC (CTCs and CTCCs). Compared to epitope-based detection platforms, the DeepPeak model demonstrated greater consistency in sensitivity compared to CellSearch, which has been reported as having anywhere between 0% to 53% sensitivity for CTCCs. Against other flow cytometer platforms, the DeepPeak model demonstrated greater sensitivity compared to the Virtual freezing fluorescence imaging flow cytometer (VIFFI FC) and comparable levels of specificity without the use of fluorescence. Finally, despite a higher FAR compared to the diffuse in vivo flow cytometer (DiFC), events detected by the DeepPeak model demonstrated higher PCC compared to the DiFC. A higher FAR was expected due to increased background signals from light scatter compared to fluorescence signals used by the DiFC for detection. In DiFC fluorescence detection, autofluorescence was the principal source of background signal and was less prevalent compared to light scatter signal.

A limitation of the DeepPeak model compared to other label-free detection platforms was the lower detection sensitivity. Higher sensitivity was achievable by reducing the number of ensemble models but came at the cost of purity, specificity, PCC, and FAR (Figure 6). Higher sensitivity would reduce the volume of blood needed to be interrogated, but poor PCC and high FAR represented undesirable artifacts in rare event detection. As such, we prioritized lower FAR and higher PCC compared to maximizing the detection sensitivity (Table 2).

**Figure 6:**
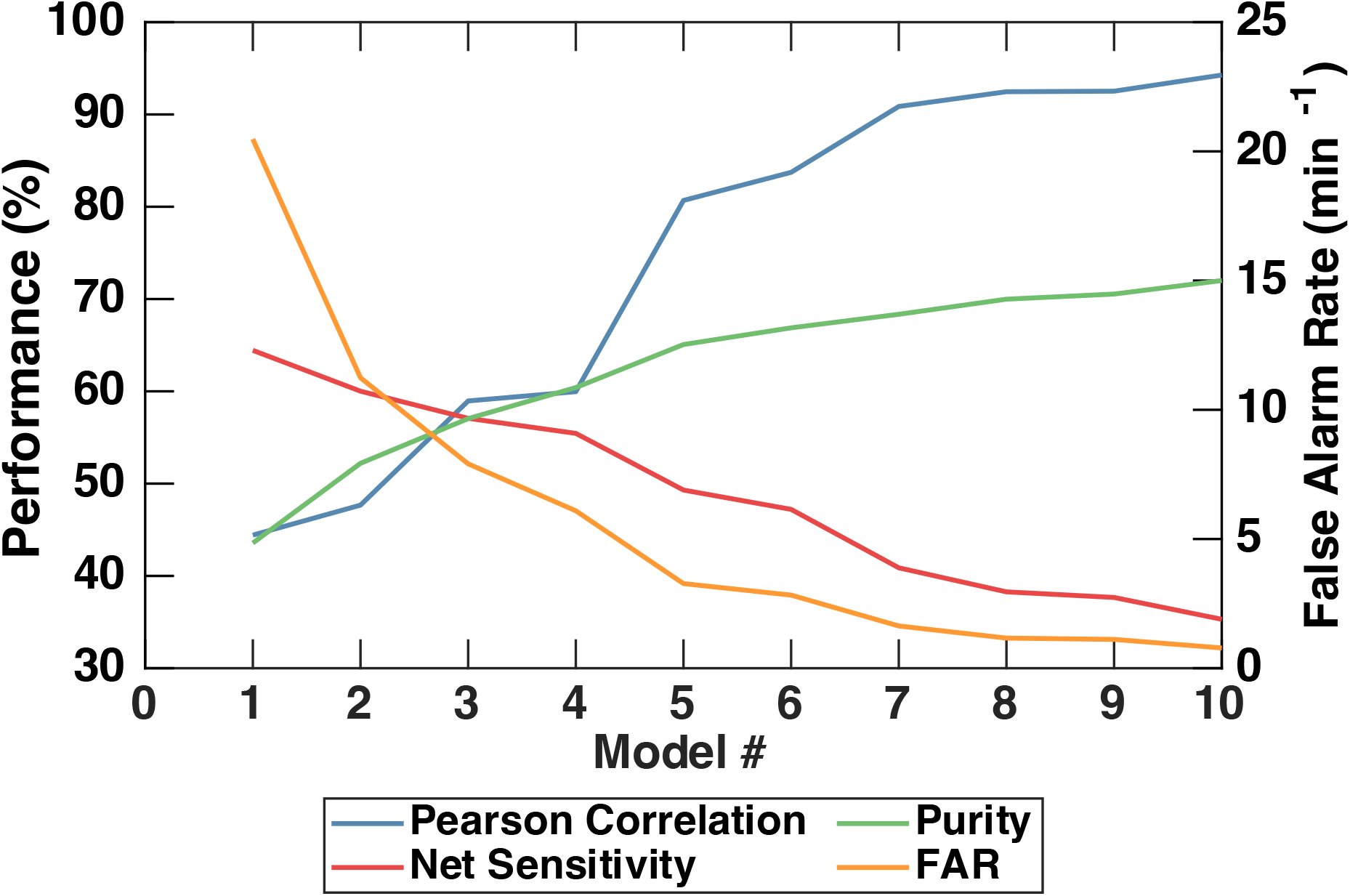
Visual summary of the impact of varying the number of ensemble models used in the ROI Classification algorithm on the DeepPeak model performance. FAR is shown on the right axis. A tradeoff between sensitivity and purity leads to changes in FAR and PCC.

**Table 2:** Comparison of all metrics for varying ensemble model counts. Numerical results correspond to numbers from **Figure 6.**

| DeepPeak Model Count | Purity | Sensitivity | Specificity | FAR | F<sub>1</sub> Score | PCC |
| --- | --- | --- | --- | --- | --- | --- |
| Single Model | 43.6% | 64.4% | 99.8% | 20.5 per minute | 0.52 | 0.44 |
| Three Models | 57.0% | 57.1% | 99.88% | 7.91 per minute | 0.57 | 0.59 |
| Ten Models | 72.0% | 35.3% | 99.97% | 0.783 per minute | 0.474 | 0.94 |

## Conclusions

In summary, we demonstrate a robust platform for label-free detection and enumeration of rare cellular events in whole blood. BSFC, in combination with deep learning models (the DeepPeak model), have implications for clinical detection and continuous monitoring of rare cellular events *in vivo*. In this study, we cover *in vitro*-based assessment of CTCC detection using the DeepPeak model. However, *in vivo* translation of the model is possible and remains the aim of our work.

Label-free detection methods face fewer regulatory barriers for clinical application than epitope-based detection methods. The DeepPeak model builds on our previous work in label-free detection of CTCCs using BSFC^12^ by implementing a more advanced signal processing algorithm for completely label-free detection of both homotypic and heterotypic CTCCs with a minimum cluster size of 2 cells. The measured FAR of 0.783 events per minute suggests that the DeepPeak model detects less than 1 FP event per every 15.2 million cellular events. To the best of our knowledge, this is the first time that FAR has been assessed for label-free CTCC detection platforms. Detected events by the DeepPeak algorithm display a high correlation with the actual number of events present within the sample despite the lower sensitivity compared to other CTCC detection platforms. The high PCC between the detected and spiked events suggests a linear relationship could be used to estimate the concentration of CTCCs from the classified events. We believe the high correlation between detected events and spike count indicates the model’s utility for extremely rare event detection. Our performance demonstrates that the DeepPeak model could be used to predict CTCC counts despite possible FP events being detected.

Label-free detection provides an inherent advantage compared to fluorescence-based methods of CTCC detection for clinical use. While the advent of new molecular probes for *in vivo* staining of CTCs and CTCCs could enable fluorescence-based *in vivo* monitoring of CTCs/CTCCs, these probes still face limitations in technical development and regulatory approval^34^. The chief advantage of BSFC and the DeepPeak model over other label-free systems is its potentially broad application to all types of cancer cell clusters *in vivo*. *In vitro* microfluidic devices enable only small volumes of blood to be sampled compared to the total peripheral blood volume. As CTC and CTCC concentrations fluctuate over time, often within the course of a couple of hours, detection of rare events in small blood volumes may lead to over or underestimation of CTCC events^24, 25^. The over or underestimation of CTCCs events could lead to poor correlation with prognosis. *In vivo* PAFC accounts for this by providing label-free, continuous *in vivo* monitoring of rare cellular events with high sensitivity and specificity. While PAFC-enabled detection of CTCs and CTCCs, it is limited to melanoma CTC/CTCC detection until probes for photoacoustic contrast are approved^31, 54–56^. These probes would face similar technical development and regulatory limitations as fluorescence-based probes preventing broad label-free monitoring of CTCCs *in vivo*.

In this study, we show that BSFC yields 72% detection purity, 99.97% net specificity, and 35.3% net sensitivity for CTCC detection. Based on this performance, 5-7 mL of blood would need to be interrogated for BSFC to detect a single CTCC. While this volume of blood is high, assessment of CTCC concentration *in vivo* could vastly impact the needed volume. Multiple studies have published conflicting concentrations of CTCCs in blood ranging from 0.44 CTCCs/mL to 10 CTCCs/mL of blood^7, 48, 49^. A challenge in assessing CTCC concentration is that all measurements to date have been collected *ex vivo* and are subject to over or underestimation. Defining an average concentration of 10 CTCCs/mL, for example, would only necessitate processing 300 μL of blood compared to 5-7 mL of blood. The range of uncertainty between concentrations highlights the need for *in vivo* detection methods to ascertain the actual concentration of CTCCs in whole blood. Here, we assume CTCC concentration is near the lower end of literature concentrations (0.4-0.5 CTCCs/ mL) to determine the maximum volume and collection time needed in a clinical setting. While processing 5-7 mL of blood is within the normal ranges for blood processing, at the throughput used in this study, processing time would approach close to 39 hours. Enhanced throughput is needed to reduce the processing time.

Multichannel flow could be used to improve BSFC throughput. A limitation in multichannel illumination and detection in our current set up is the available slit characteristics (5 x 30 μm^2^). Future efforts will be focused on implementing straightforward modifications to our illumination scheme and microfluidic device design to enable data collection from whole blood flowing through multiple microfluidic channels.

To understand the limitations of detection sensitivity, we carefully examined FN peaks from the ROI Classification algorithm (Sample FN peaks are included in Supplementary Fig. S4 (online)). Considering the FWHM of all mislabeled events, >80% of mislabeled events were 2-cell CTCCs or potentially 2-cell CTC-WBC events, 14-16% were 3-6 cell CTCCs or 3-6 cell CTC-WBC events, and less than 4% of mislabeled events were 6+ cell CTCCs or 6+ cell CTC-WBC events. This suggests that the classification errors center around mostly smaller CTCC events, which are known to be more challenging to classify from large single-cell events and WBCs. While these events could be excluded to achieve improved performance, these events were included as the role of smaller clusters may be significant. Intriguingly, detection purity remained constant between the test set and the full dataset while detection sensitivity decreased. This would suggest that the classification model has sufficiently learned parameters for FPs but was limited by the number of TP (CTCC) peaks included in training. The disparity in CTCC (16,243) and NC peaks (286,618) in the training set likely accounts for the difference between training and test set performance. A greater distribution of CTCC data in the training data could improve sensitivity. As we move forward, we aim to collect more training data and examine alternative training schemes to prioritize maximizing sensitivity, such as implementing data augmentation.

In our prior work, we examined the limit in our classification performance using a simplified system with fewer illumination parameters^12^. Detection with two or even one illumination wavelength can vastly reduce data complexity, instrumentation cost, and improve the adoption of portable systems. We previously reported that detection based on only two interrogation wavelengths was sufficient to achieve comparable levels of performance as using all three interrogation wavelengths^12^. In this study, we observed similar results when using only two of the three interrogation wavelengths for classification (see Supplementary Fig. S5 online). Surprisingly, using only 405 nm excitation, also led to comparable levels of classification sensitivity and PCC, albeit with a loss in detection purity. This suggests that sensitivity was highly correlated with the 405 nm channel. Further exploration is needed to determine how optimization of the 488 and 633 nm channels data could improve model sensitivity.

*In vivo* translation of BSFC with the DeepPeak model is yet to be explored. However, as a label-free detection and monitoring platform, clinical translation of BSFC holds promise. The use of deep learning is instrumental in the accurate and sensitive detection of CTCCs in noisy, label-free blood scatter data. As a greater number of datasets and optimization of instrumental setup becomes available, we plan to present a more advanced BSFC capable of sensitive detection of CTCCs in whole blood with high throughput. In addition to *in vitro* throughput enhancement, we aim to address *in vivo* throughput, addressing one of the major limitations of label-free microscopy-based flow cytometry^5, 34^. In its current form, label-free BSFC has potential uses in the non-destructive isolation of CTCCs, *ex vivo* monitoring of CTCC dynamics, and *ex vivo* treatment monitoring. However, we aim to demonstrate expanded uses of BSFC in the near future through *in vivo* detection of CTCCs in whole blood.

## Statement of Contributions

N.V., under guidance by I.G., conducted all experiments, analyzed data, and prepared all figures. With guidance from I.G., A.P., and P.S., N.V. developed the DeepPeak model. M.E. aided with all animal-related work and acquisition of all blood. I.G. supervised the project and along with N.V. prepared the manuscript text. All authors have reviewed and approved the manuscript.

## Acknowledgements

We would like to thank the National Institute of Biomedical Imaging and Bioengineering (R03 EB027363) and National Cancer Institute (R21 CA271679) for funding this work. We would also like to thank Dr. Madeleine Oudin (Tufts University) for her guidance with cell culture and providing us with the necessary cell lines for this study. The authors acknowledge the Tufts University High Performance Compute Cluster (https://it.tufts.edu/high-performance-computing) which was utilized for the research reported in this paper. Finally, we would like to thank Dr. Jeffrey Guasto (Tufts University) for their guidance on microfluidic design and device fabrication and Dr. Shannon Stott (Massachusetts General Hospital) for her guidance on interpreting results in the field of CTCC detection.

