## Supplementary for "Deep Learning-Enabled, Detection of Rare Circulating Tumor Cell Clusters in Whole Blood Using Label-free, Flow Cytometry"

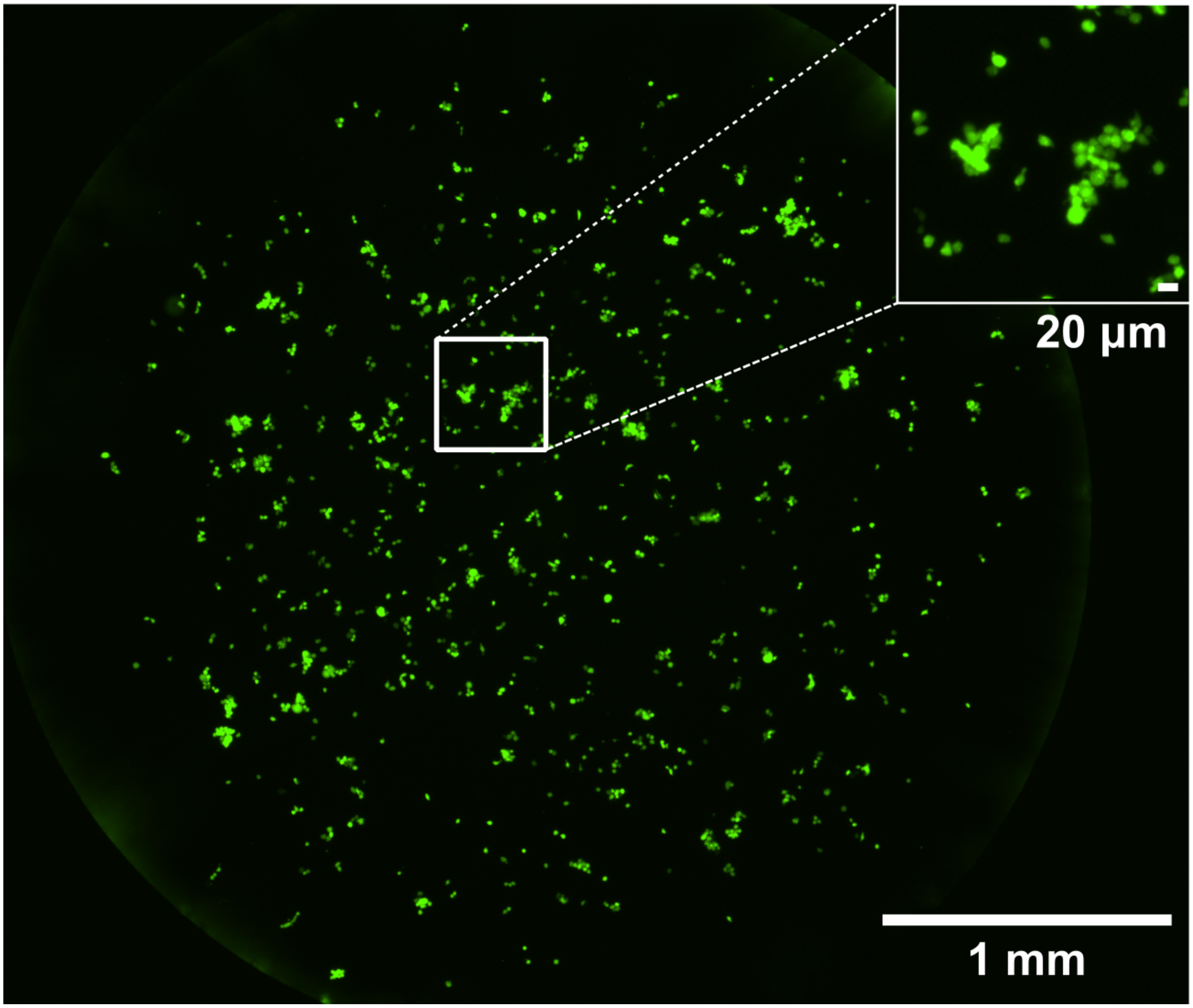


Supplementary Figure S1: Fluorescence image of CTCCs generated prior to spiking in whole blood. CTCs and CTCCs are observed in the spiked samples. CTCCs range from 2-9+ cells in size. Large single cells (~20 μm) are also observed in the image. On average, CTCs are 13 μm in size. The image shown was acquired using a Nikon Eclipse Ti2 with a Plan Fluor 4x PhL DL objective (NA =0.13).

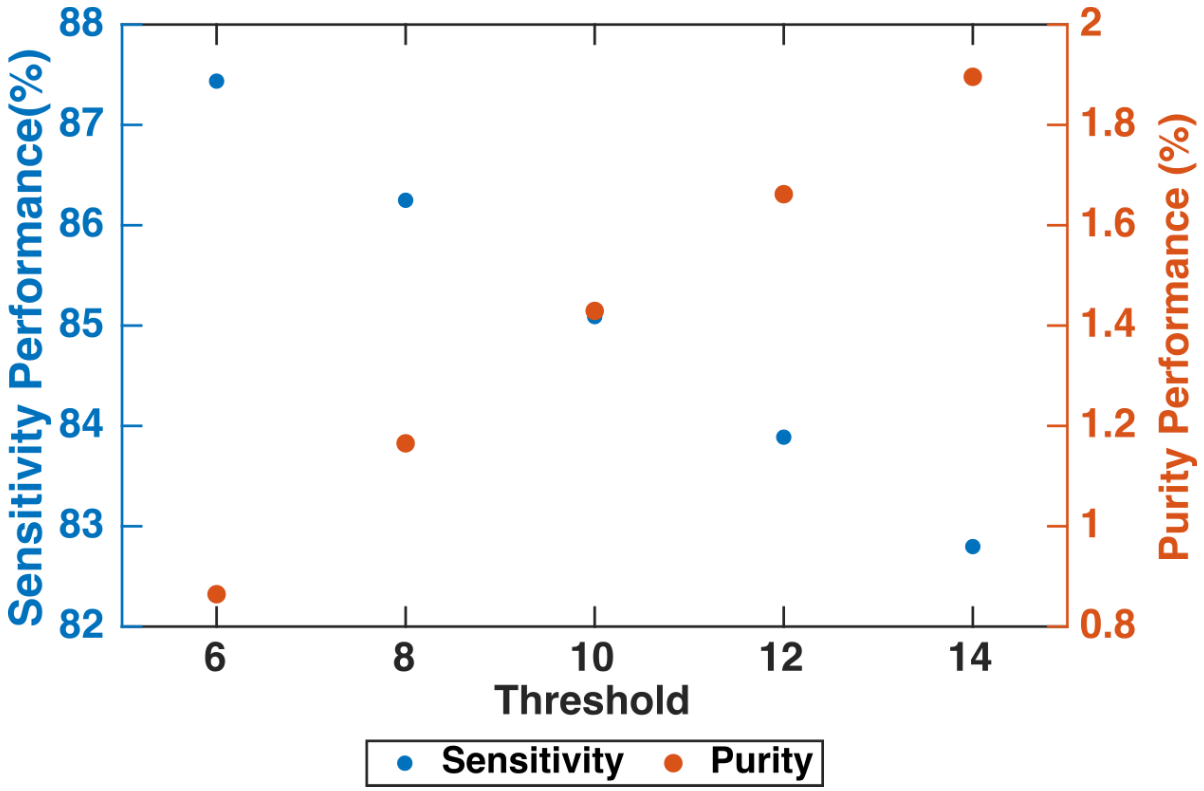


Supplementary Figure S2: Variations in the intensity threshold used by the ROI Detection Algorithm impact the detection sensitivity and purity inversely. A threshold of 10 was selected to maximize both sensitivity and purity.

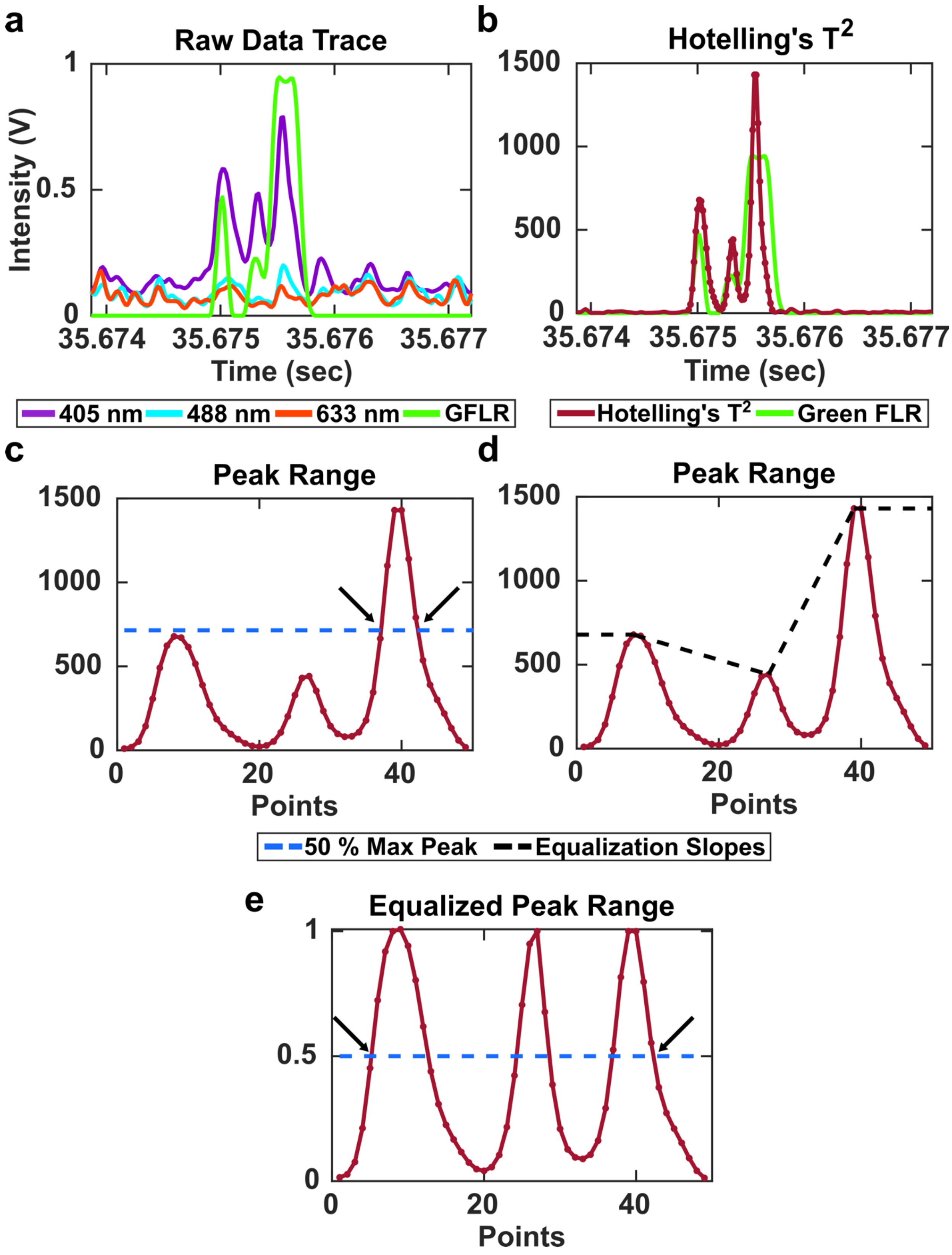


Supplementary Figure S3: **(a)** Raw data trace for a CTCC peak with weak shoulder peaks. **(b)** After PCA and Hotelling's $T^2$ test, the weak shoulder peaks are still present.

**(c)** Using our standard intensity threshold algorithm to define the range of points within the cluster, 49 points are selected. The half maximum is calculated (drawn in blue) and full-width-at-half-max (FWHM) is calculated based on the left and rightmost points where the signal crosses the half maximum (indicated with black arrows). This leads to a detected FWHM of 6 points (Corresponding to a narrow single-cell event). **(d)** We use peak equalization to correct for the incorrectly detected narrow peak, calculating the slopes between each peak and normalizing the data based on these slopes. **(e)** The equalized peaks are scaled between zero to one with a half maximum = 0.5 (blue line). The FWHM is again assessed (black arrows) and is now 38 points, correctly indicating a 3-6 cell CTCC event.

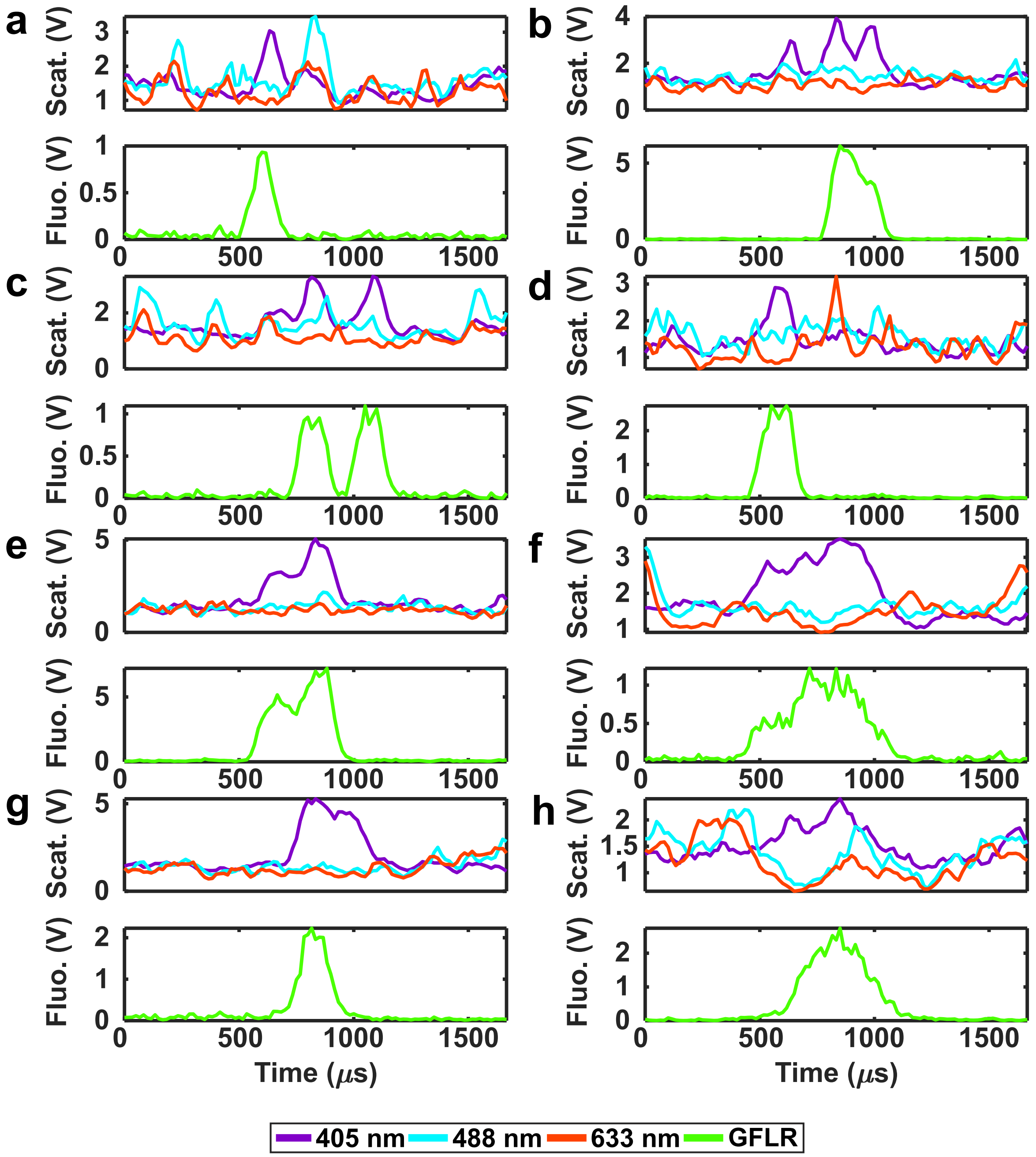


Supplementary Figure S4: Sample raw traces of false negative peaks from the ROI Classification algorithm. False negative events include **(a,d,g)** a 2-cell heterogenous CTC-WBC event, **(b,c)** 3-6 cell heterogenous CTCC, **(e,h)** a homogenous 2-cell CTCC, and **(f)** a 3-6 cell homogenous CTCC. All labels are defined based on FWHM.

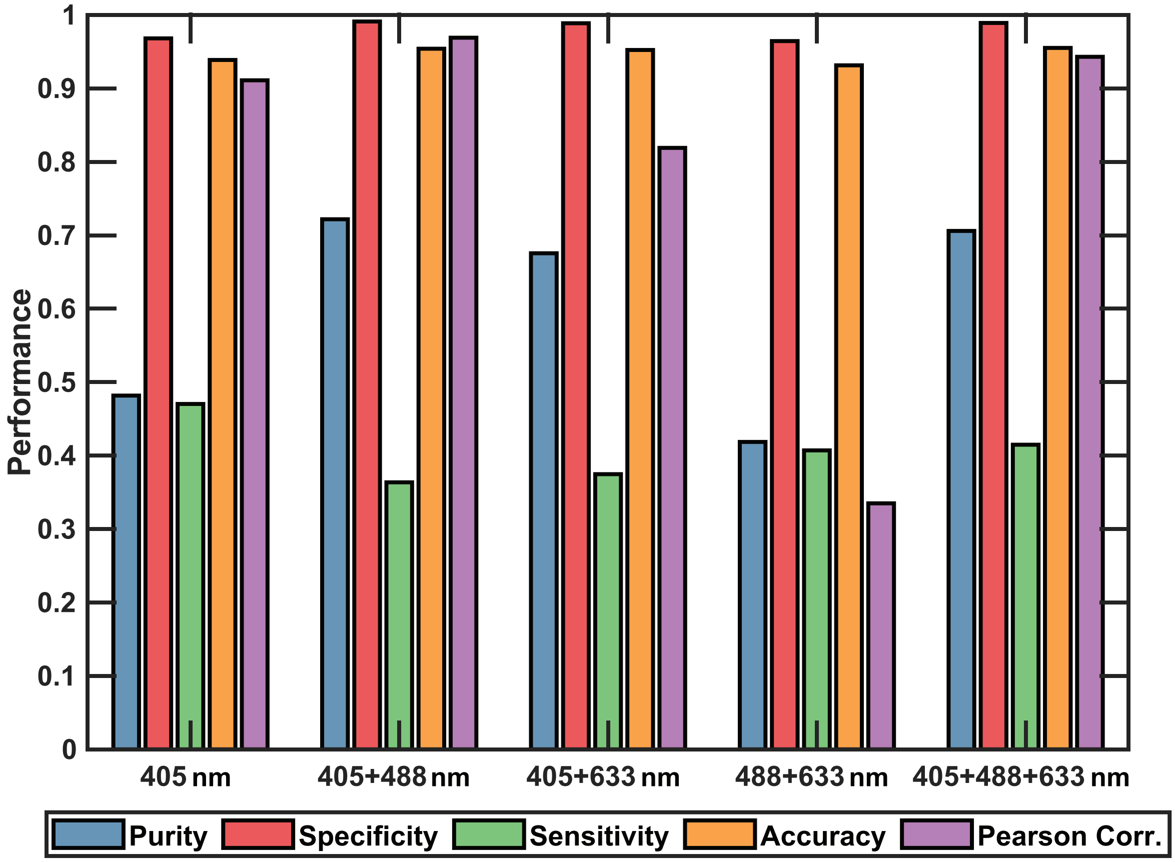


Supplementary Figure S5: Simplified datasets were trained and evaluated using an identical model architecture. Interrogation wavelengths were permuted to identify which laser sources were needed to achieve desirable performance. We observe 405 + 488 or 405 + 633 can achieve similar performance as 405 + 488 + 633. Further, using 405 only could also achieve comparable performance for all metrics outside of detection purity.
